# Secondary metabolites produced during *Aspergillus fumigatus* and *Pseudomonas aeruginosa* biofilm formation

**DOI:** 10.1101/2022.04.25.489484

**Authors:** Rafael Wesley Bastos, Daniel Akiyama, Thaila Fernanda dos Reis, Ana Cristina Colabardini, Rafael Sanchez Luperini, Patrícia Alves de Castro, Regina Lúcia Baldini, Taícia Fill, Gustavo H. Goldman

**Affiliations:** Faculdade de Ciências Farmacêuticas de Ribeirão Preto, Universidade de São Paulo, Brazil; Instituto de Química, Universidade Estadual de Campinas, Brazil; Departamento de Bioquímica, Instituto de Química, Universidade de São Paulo, Brazil

## Abstract

In Cystic Fibrosis (CF), mucus plaques are formed in the patient’s lung, creating a hypoxic condition and a propitious environment for colonization and persistence of many microorganisms. There is clinical evidence showing that *Aspergillus fumigatus* can co-colonize CF patients with *Pseudomonas aeruginosa*, which has been associated with lung function decline. *P. aeruginosa* produces several compounds with inhibitory and anti-biofilm effects against *A. fumigatus* in vitro; however, little is known about the fungal compounds produced in counterattack. Here, we annotated fungal and bacterial secondary metabolites (SM) produced in mixed biofilms in normoxia and hypoxia conditions. We detected nine SMs produced by *P. aeruginosa*. Phenazines and different analogs of pyoverdin were the main compounds produced by *P. aeruginosa*, and their secretion were increased by the fungal presence. The roles of the two operons responsible for phenazines production (*phzA1* and *phzA2*) were also investigated showing both mutants are able to produce partial sets of phenazines. We detected a total of 20 SMs secreted by *A. fumigatus* either in monoculture or in co-culture with *P. aeruginosa*. All these compounds are secreted during biofilm formation either in normoxia or hypoxia. However, only eight compounds (demethoxyfumitremorgin C, fumitremorgin, ferrichrome, ferricrocin, tricetylfusigen, gliotoxin, gliotoxin E, and pyripyropene A) were detected during the biofilm formation by the co-culture of *A. fumigatus* and *P. aeruginosa* upon both normoxia and hypoxia conditions. Overall, we showed how diverse is SM secretion during *A. fumigatus* and *P. aeruginosa* mixed culture and how this can affect biofilm formation both in normoxia and hypoxia.

**Author Summary:** The interaction between *Pseudomonas aeruginosa* and *Aspergillus fumigatus* has been well-characterized *in vitro*. In this scenario, the bacterium exerts a strong inhibitory effect against the fungus. However, little is known about the metabolites produced by the fungus to counterattack the bacteria. Our work aimed to annotate secondary metabolites (SM) secreted during co-culture between *P. aeruginosa* and *A. fumigatus* during biofilm formation in both normoxia and hypoxia. The bacterium produces several different types of phenazines and pyoverdins, in response to the fungus presence. In contrast, we were able to annotate 29 metabolites produced during *A. fumigatus* biofilm formation but only eight compounds were detected during biofilm formation by the co-culture of *A. fumigatus* and *P. aeruginosa* upon both normoxia and hypoxia. In conclusion, we have detected many SMs secreted during *A. fumigatus* and *P. aeruginosa* biofilm formation. This analysis can provide several opportunities to understand the interaction between these two species.

## Introduction

*Pseudomonas aeruginosa* is a gram-negative bacterium that grows aerobically and in anaerobic conditions under certain specific circumstances. This species is ubiquitous in nature and has been found inhabiting soil and water, and colonizing humans, sometimes acting as an opportunistic pathogen (1). In immunosuppressed, burnt, and hospitalized patients, *P. aeruginosa* is responsible for a broad spectrum of serious diseases ranging from acute to chronic infections, such as bloodstream infections in intensive care units (ICU), surgical site infections, hospital-acquired pneumonia, respiratory and urinary tract infections, and burn and chronic dermal wound infections (1, 2). *P. aeruginosa* also chronically infects the lungs of people with underlying pulmonary diseases, as cystic fibrosis (CF).

CF is a genetic disorder caused by mutations in CF transmembrane conductance regulator (CFTR) gene resulting in defective chloride secretion, altered airway surface liquid, ciliary dyskinesis and impaired mucociliary clearance (3). Such changes lead to motionless mucus plaques, which creates a hypoxic condition and a propitious environment to colonization and persistence of many microorganisms, notably *P. aeruginosa* (1, 4). By performing an *in vivo* characterization of CF airways, Worlitzsch and colleagues (4) demonstrated that *P. aeruginosa* forms biofilm-like macrocolonies in the intraluminal site, which is markedly hypoxic due to mucus accumulation. In response to hypoxia, *P. aeruginosa* increases alginate exopolysaccharide production, and that may help the bacteria grow as biofilm and persist in that environment. In fact, biofilm formation is a fundamental factor for *P. aeruginosa* persistence in chronic infections because it confers increased resistance to antimicrobials and immune killing, which explains why it is virtually impossible to eradicate such infections (5, 6).

In addition to *P. aeruginosa* and other bacteria, CF lungs can be colonized by fungi, being *Aspergillus fumigatus* the main isolated mold. There is a particular interest in this opportunistic pathogen as *A. fumigatus* presence in respiratory CF samples has been associated with poorer prognosis and pulmonary function decline (7). *A. fumigatus* is a ubiquitous filamentous fungus encountered in soil, water, air, decomposing organic matter and plant-based materials (8, 9) and it probably evolved in contact with water and soil bacteria as *P. aeruginosa*. *A, fumigatus* can cause a range of illnesses varying from chronic or hypersensitization (allergic reactions) disorders to invasive and life-threatening diseases (10). In CF patients, *A. fumigatus* may colonize the bronchi, which is frequently accompanied by hypersensitization (1, 11, 12), allergic bronchopulmonary aspergillosis (ABPA) (13), and bronchitis (14).

There is clinical evidence that *A. fumigatus* and *P. aeruginosa* co-colonize CF patients, which is associated with lung function decline. Some reports estimates that 60% of patients with chronic *P. aeruginosa* infection also carry *A. fumigatus* (15, 16, 17, 18). Studies investigating how *A. fumigatus*-*P. aeruginosa* affect each other *in vivo* and the outcome of this interaction for the host are limited; however, several studies have analyzed this interaction *in vitro*. Overall, *P. aeruginosa* has a strong inhibitory effect against *A. fumigatus in vitro* (including inhibition of biofilm formation and conidiation) due to bacteria produced compounds (1, 19). The surfactant dirhamnolipids inhibit fungal growth by blocking β-1,3 glucan synthase (20), a key enzyme for cell wall production; the quorum sensing homoserine lactones act suppressing hyphal growth (21); the siderophore pyoverdine causes fungal iron starvation (19, 22, 23), and pyochelin and phenazines kill *A. fumigatus* by inducing oxidative and nitrosative stresses as well as iron starvation (19, 24).

Phenazines are nitrogen-containing colored aromatic molecules, which constitute a large group of secondary metabolites (SM) produced by bacteria with broad physiological functions, acting as antibiotics (2) and antifungals (24, 25, 26, 27), involved in biofilm formation (28), and regulation of gene expression (29). Several phenazines are produced by *P. aeruginosa*, such as pyocyanin (PYO); phenazine-1-carboxamide (PCN); phenazine-1-carboxylic acid (PCA); 1-hydroxyphenazine (1-OHphz), and 5-methyl-phenazine-1-carboxylic acid (5-Me-PCA). Biosynthesis of PCA from chorismic acid requires enzymes coded by two sets of homologous genes (*phzABCDEFG*) located in two nearly identical redundant operons (*phz1* and *phz2*), but with different promoters and flanking regions (30). Other three genes, *phzM*, *phzS* and *phzH* code for enzymes that convert PCA to PYO and PCN and are located next to either *phz1* or *phz2* operons (30).

Although *P. aeruginosa* compounds with inhibitory and anti-biofilm effects against *A. fumigatus* have been revealed, little is known about the compounds produced by the fungus during interaction. Furthermore, most of the studies about *P. aeruginosa* SMs during interaction were done by using mutants lacking important genes for biosynthesis of such compounds, and/or by measuring the effect of adding purified compounds or bacterial filtered supernatant to the co-culture or monoculture. There is a lack of studies identifying fungal and bacterial SM produced throughout co-culturing and mixed biofilms, and their roles in the fungal-bacterial interaction, especially regarding fungal metabolites. Little attention has also been given to the differences in SM production and their effects on biofilm production under hypoxia, a special atmosphere that must be faced by respiratory pathogens.

Here, we show the SM produced by *P. aeruginosa* and *A. fumigatus* in single or mixed biofilms during normoxia and hypoxia conditions. We detected ten SMs produced by *P. aeruginosa.* Phenazines and different analogs of pyoverdin were the main compounds produced by *P. aeruginosa*, and their secretion were increased in the fungal presence. The effects of the two operons that regulates phenazines production (*phzA1* and *phzA2*) was also investigated. The results showed that *phzA1* and Δ*phzA2* mutants can produce a subset of phenazines when in hypoxia and in the presence of the fungus. In contrast, we were able to detect 20 SMs produced by *A. fumigatus*, but only eight of them (demethoxyfumitremorgin C, fumitremorgin C, ferrichrome, ferricrocin, triacetylfusigen, gliotoxin, gliotoxin E, and pyripyropene A) were produced in the presence of *P. aeruginosa*.

## Results

### *A. fumigatus* and *P. aeruginosa* biofilm formation in normoxia and hypoxia

We established a protocol for *P. aeruginosa* and *A. fumigatus* biofilm formation under normoxia and hypoxia by using 10^5^ colony formation units (CFUs)/ml from exponential phase *P. aeruginosa* cultures and *A. fumigatus* 10^6^ conidia/ml (Figure 1). *P. aeruginosa* wild-type (Pa), *phzA1* (A1) and Δ*phzA2* (A2) mutant strains have comparable biofilm formation. Mixed biofilms of *A. fumigatus*-*P. aeruginosa* (AfPa), *A. fumigatus*-*P. aeruginosa phzA1* (AfA1), and *A. fumigatus*-*P. aeruginosa* Δ*phzA2* (AfA2) have more biomass than bacteria-only biofilms, in both normoxia and hypoxia (Figures 1A and 1B). However, Af biofilms without bacteria present more biomass than mixed biofilms, indicating an antagonist role of *P. aeruginosa* towards Af biofilm formation.

**Figure 1.**
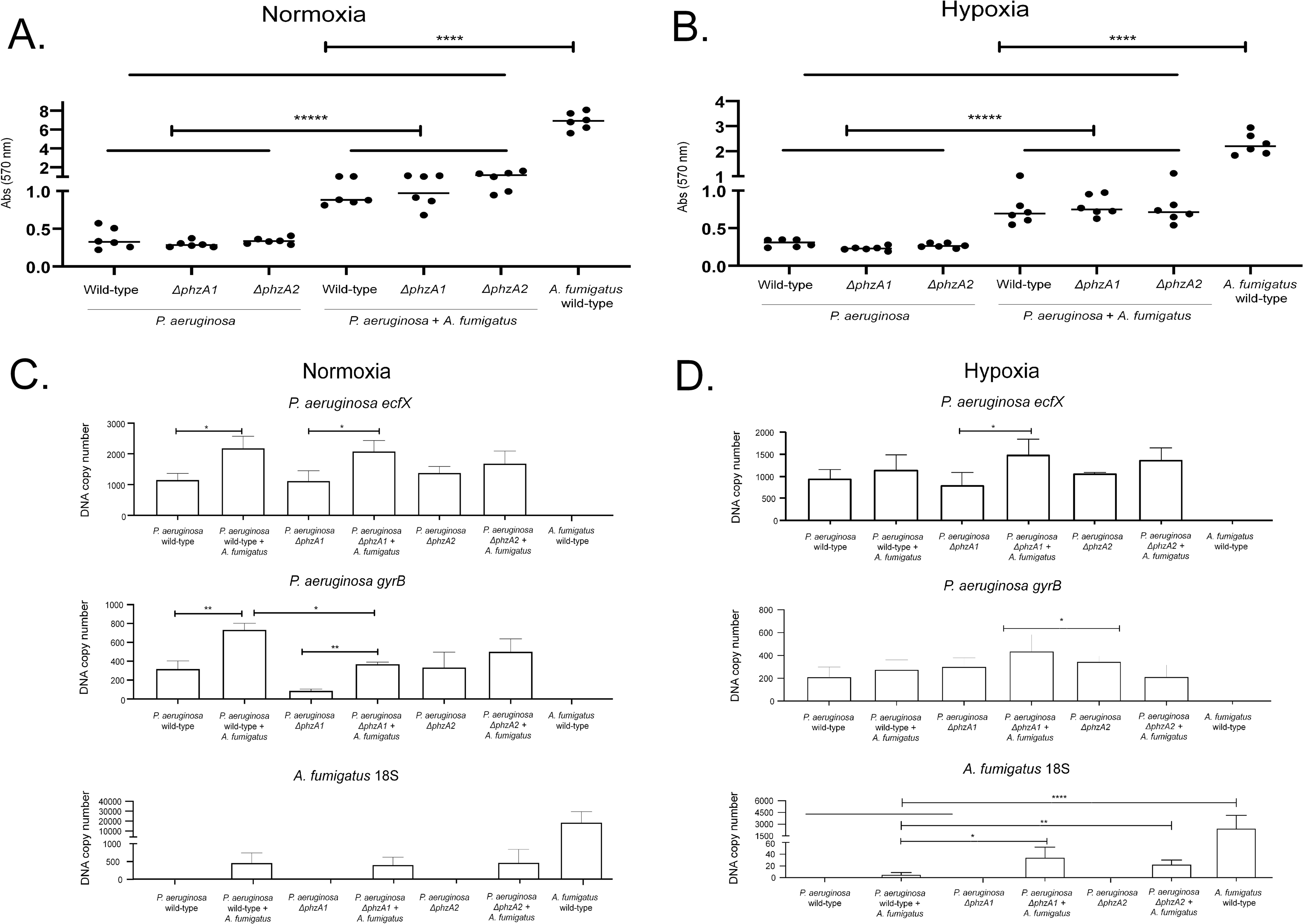
Biofilm formation by *P. aeruginosa* and *A. fumigatus*. (A) and (B) *P. aeruginosa* and *A. fumigatus* have grown for 5 days at 37 °C in normoxia and hypoxia conditions. The results of the absorbance of crystal violet are the average of six repetitions ± standard deviation. (C) and (D) qPCR for *P. aeruginosa efcX* and *gyrB*, and *A. fumigatus* 18S. The resuls are the average of three repetitions ± standard deviation.

These results were refined by estimating the *P. aeruginosa* and *A. fumigatus* DNA copy number by qPCR using *P. aeruginosa ecfX* (encoding an extracytoplasmic function sigma factor unique to *P. aeruginosa,* also annotated as *hxuI*) and *gyrB* (encoding a DNA gyrase), and *A. fumigatus* 18S DNA (31, 32, 33). Corroborating the biofilm biomass results, Af DNA copy number decreases in the presence of any *P. aeruginosa* strain in all conditions (Fig 1C and1D). In normoxia conditions, we observed that *P. aeruginosa* Pa and A1 DNA copy number increases in the presence of Af, but this is not true for A2, that show the same DNA copy number with or without Af (Figure 1C). In contrast, in hypoxia conditions, only AfA1 shows a higher *P. aeruginosa* DNA copy number than A1 alone (Figure 1D); interestingly, there is increased *A. fumigatus* DNA copy number in AfA1 and AfA2 as compared to AfPa (Figure 1D). This suggests that *P. aeruginosa phz* mutant strains have a lower ability to inhibit *A. fumigatus* biofilm than the wild type strain.

These results strongly indicate that we have established a robust *A. fumigatus* and *P. aeruginosa* biofilm formation protocol under normoxia and hypoxia conditions. Next, we used HPLC-HRMS^2^ to identify SMs in the *A. fumigatus* and *P. aeruginosa* biofilm supernatants. We were able to annotate a total of 29 SMs, 9 from *P. aeruginosa* and 20 from *A. fumigatus* in the supernatants produced in any condition (Supplementary Table S1, Supplementary Figures S1 to S8 at <u>10.6084/m9.figshare.19620702</u> and Figure 2).

**Figure 2.**
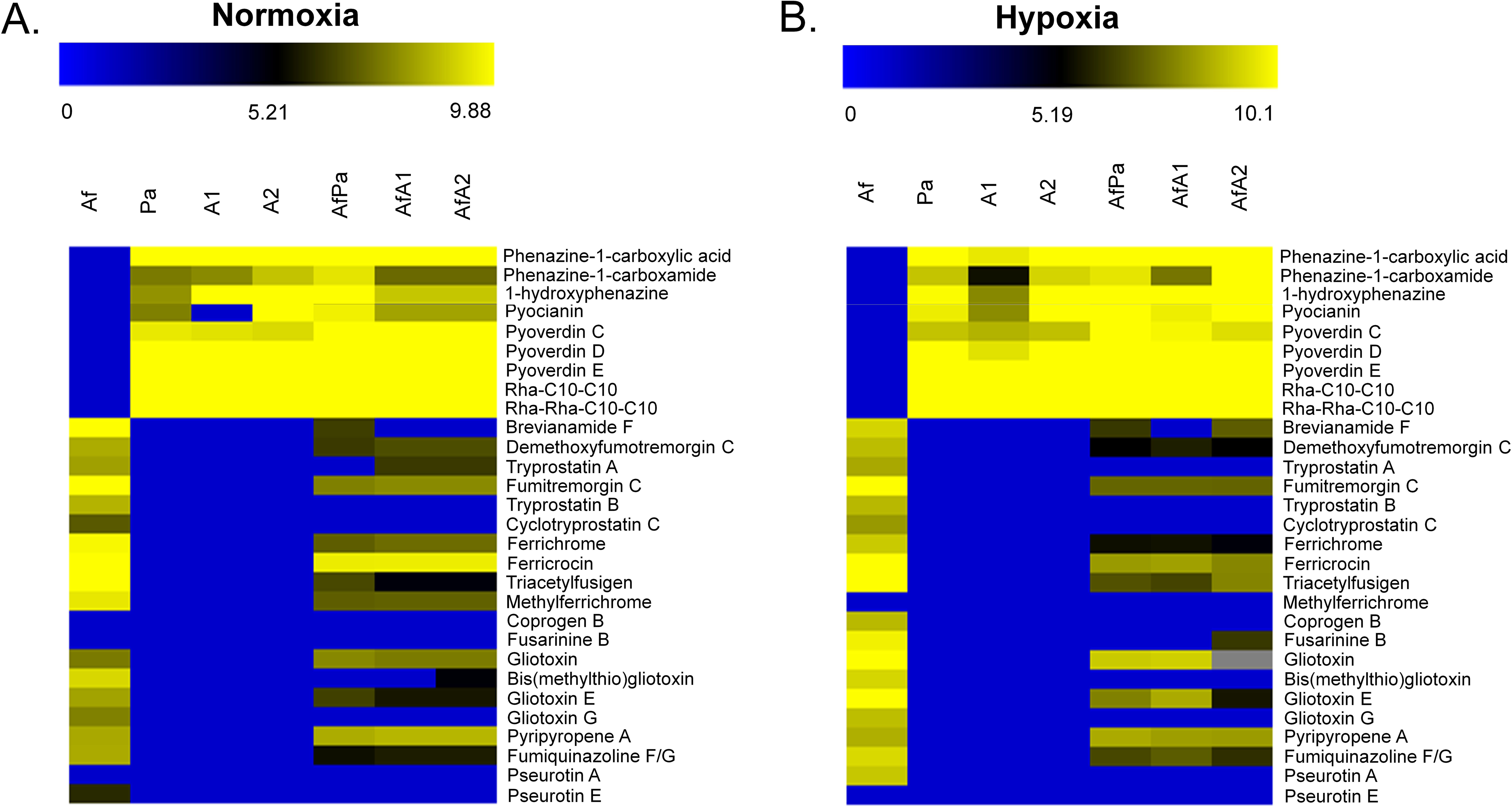
Specialized metabolite (SM) production by *P. aeruginosa* and *A. fumigatus*. (A) and (B) Heat maps depicting the log10 of the area of chromatograms of 29 SMs (9 from *P. aeruginosa* and 20 from *A. fumigatus*). The values represent the average of three independent biological repetitions.

### Secondary metabolites produced by *P. aeruginosa* during biofilm formation

PhZA1-G1 and PhZA2-G2 pathways use chorismic acid as the precursor for transformation into phenazine-1-carboxylic acid (PCA) (Figure 3A). PCA is converted into phenazine-1-carboxamide, 1-hydroxiphenazine, and 5-methylphenazine-1-carboxylic acid betaine by PhzH, PhzS, and PhzM, respectively (Figure 3A). Subsequently, 5-methylphenazine-1-carboxylic acid betaine is converted into pyocyanin by PhzS (Figure 3A). As expected, methylphenazine-1-carboxylic acid betaine and pyocyanin were not detected in the *phzA1* mutant (A1 biofilm) under normoxia conditions (Figures 2, 3C and 3E). All these compounds except methylphenazine-1-carboxylic acid betaine are induced in mixed AfPa biofim in normoxia and hypoxia conditions, as compared to Pa-only biofilm (Figures 2, 3A to 3E). Interestingly, the production of phenazines in A1 and A2 biofilms vary when compared to the wild type Pa biofilm: whereas A1 presents low phenazine production in all conditions, A2 has higher phenazines production than Pa and A1.

**Figure 3.**
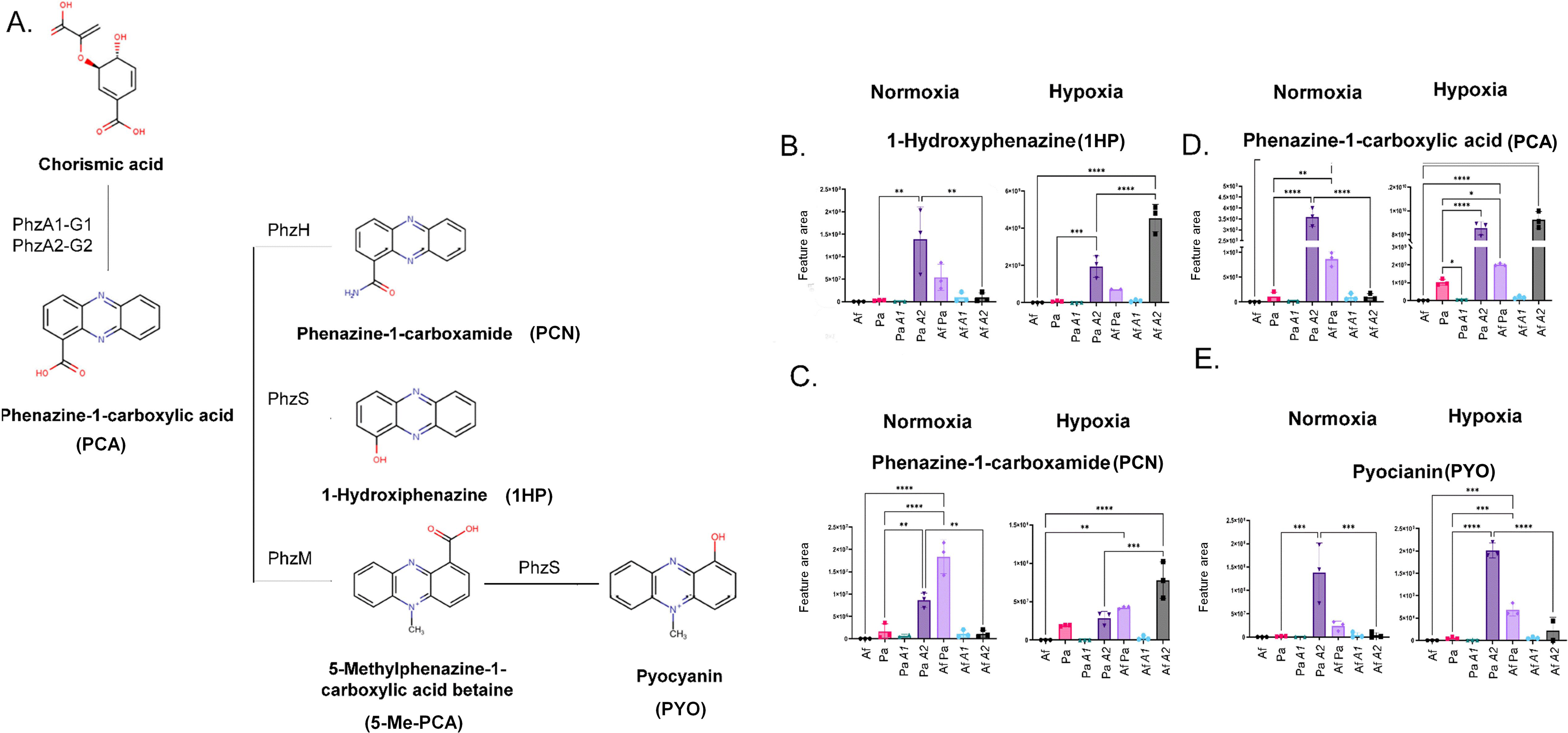
*P. aeruginosa* phenazine production during biofilm formation in normoxia and hypoxia. (A) *P. aeruginosa* phenazine biosynthesis pathway. Areas of the chromatograms of (B) 1-hydroxiphenazine, (C) phenazine-1-carboxylic acid, (D) phenazine-1-carboxamide, and (E) pyocianin. The results are the average of three repetitions ± standard deviation. Pa = *P. aeruginosa* wild-type; Pa*A1 = P. aeruginosa phzA1* mutant; PaA2 = *P. aeruginosa ΔphzA2* mutant; AfPa = *A. fumigatus* + *P. aeruginosa* wild-type; Af*A1 = A. fumigatus + P. aeruginosa phzA1* mutant; and AfA2 = *A. fumigatus* + *P. aeruginosa ΔphzA2* mutant.

When phenazine production by AfA1 mixed biofilms are analyzed, there is an increase for all compounds both in normoxia and hypoxia conditions, as compared to A1-only biofilm; however, AfA2 biofilms only increase phenazine production in hypoxia, as compared to A2-only biofilms, except for PYO. Phenazine-1-carboxamide production has comparable amounts in the *P. aeruginosa* wild-type Pa and A1 strains in hypoxia conditions (Figures 2B, 3A to 3E). Phenazine production in AfA1 is much lower than AfPa in both conditions (Figures 2, 3A to 3E). As expected, phenazine production is significantly higher during biofilm formation in hypoxia than normoxia (Figures 2, 3A to 3F), as PYO can be used as an alternative electron acceptor in anaerobiosis (34).

The siderophores pyoverdine C, D, and E and Rha-C10-C10 and Rha-Rha-C10-C10 ramnolipids are produced in comparable amounts during Pa, A1, and A2 biofilm formation under normoxia and hypoxia conditions (Figures 2 and 4). The *phzA1* and Δ*phzA2* mutations affect the pyoverdin D and E during biofilm formation in hypoxia (Figures 2B, 4B and 4C). All these compounds are produced in larger amounts in mixed biofilms (AfPa, AfA1 and AfA2) in both normoxia and hypoxia, except for pyoverdine E under normoxia (Figures 2, 4A to 4E).

**Figure 4.**
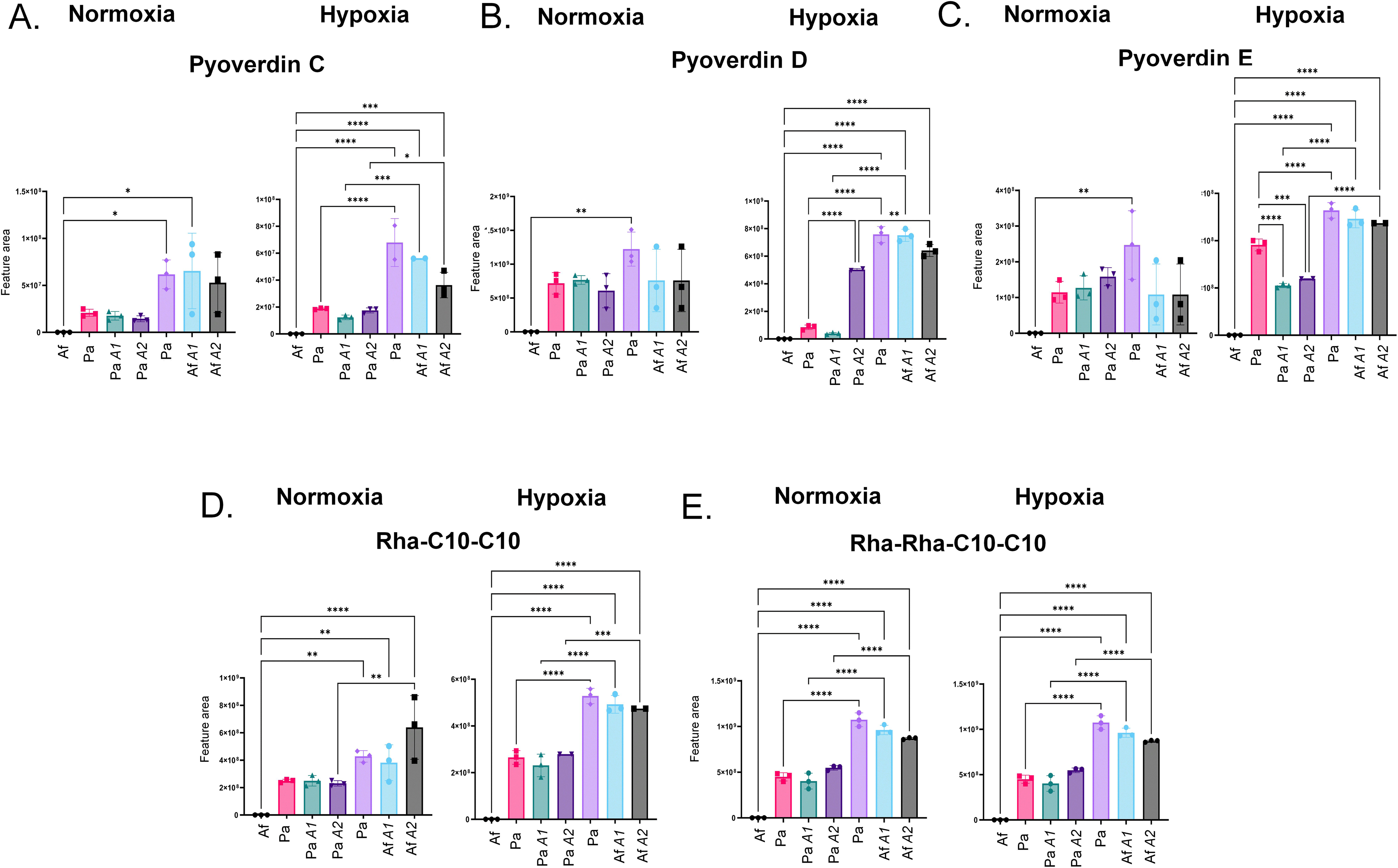
*P. aeruginosa* pyoverdin and rhamnolipid production during biofilm formation in normoxia and hypoxia. Areas of the chromatograms of (A) pyoverdin C, (B) pyoverdin D, (C) pyoverdin E, (D) Rha-C10-C10, and (F) Rha-Rha-C10-C10. The results are the average of three repetitions ± standard deviation. Pa = *P. aeruginosa* wild-type; Pa*A1 = P. aeruginosa phzA1* mutant; PaA2 = *P. aeruginosa ΔphzA2* mutant; AfPa = *A. fumigatus* + *P. aeruginosa* wild-type; Af*A1 = A. fumigatus + P. aeruginosa phzA1* mutant; and AfA2 = *A. fumigatus* + *P. aeruginosa ΔphzA2* mutant.

These results indicate that interaction with Af in mixed biofilms stimulates the production of phenazines by the Pa WT strain, and that mutation in *phzA1* or *phzA2* modulates negative or positively phenazine production, respectively. Pyoverdine and rhamnolipid production are not significantly different among Pa, A1, and A2 but increased production is detected in all mixed biofilms (AfPa, AfA1, and AfA2).

### *A. fumigatus* biofilm formation induces the production of metabolites in the superpathway of fumitremorgin biosynthesis

The tremorgenic mycotoxins fumitremorgins are a group of prenylated indole alkaloids produced by *A. fumigatus* (35). Fumitremorgin C is produced through a series of steps in the superpathway of fumitremorgin biosynthesis (http://vm-trypanocyc.toulouse.inra.fr/META/NEW-IMAGE?type=PATHWAY&object=PWY-7525&orgids=LEISH; Figure 5A). Several metabolites in this pathway are produced during biofilm formation under normoxia and hypoxia conditions, such as brevianamide F, demethoxyfumitremorgin C, tryprostatin A, fumitremorgin C, tryprostatin B, and cyclotryprostatin (Figures 2, 5B to 5G). Although lower than Af, AfPa biofilm shows production of brevianamide F (in hypoxia), demethoxyfumitremorgin C (normoxia and hypoxia), and fumitremorgin C (normoxia and hypoxia); there are no differences between AfPa, AfA1 and Af2 (Figures 2, 5B, 5E, and 5F). These results suggest that *P. aeruginosa* is able to inhibit completely or partially the production of several metabolites in the superpathway of fumitremorgin biosynthesis during AfPa biofilm formation.

**Figure 5.**
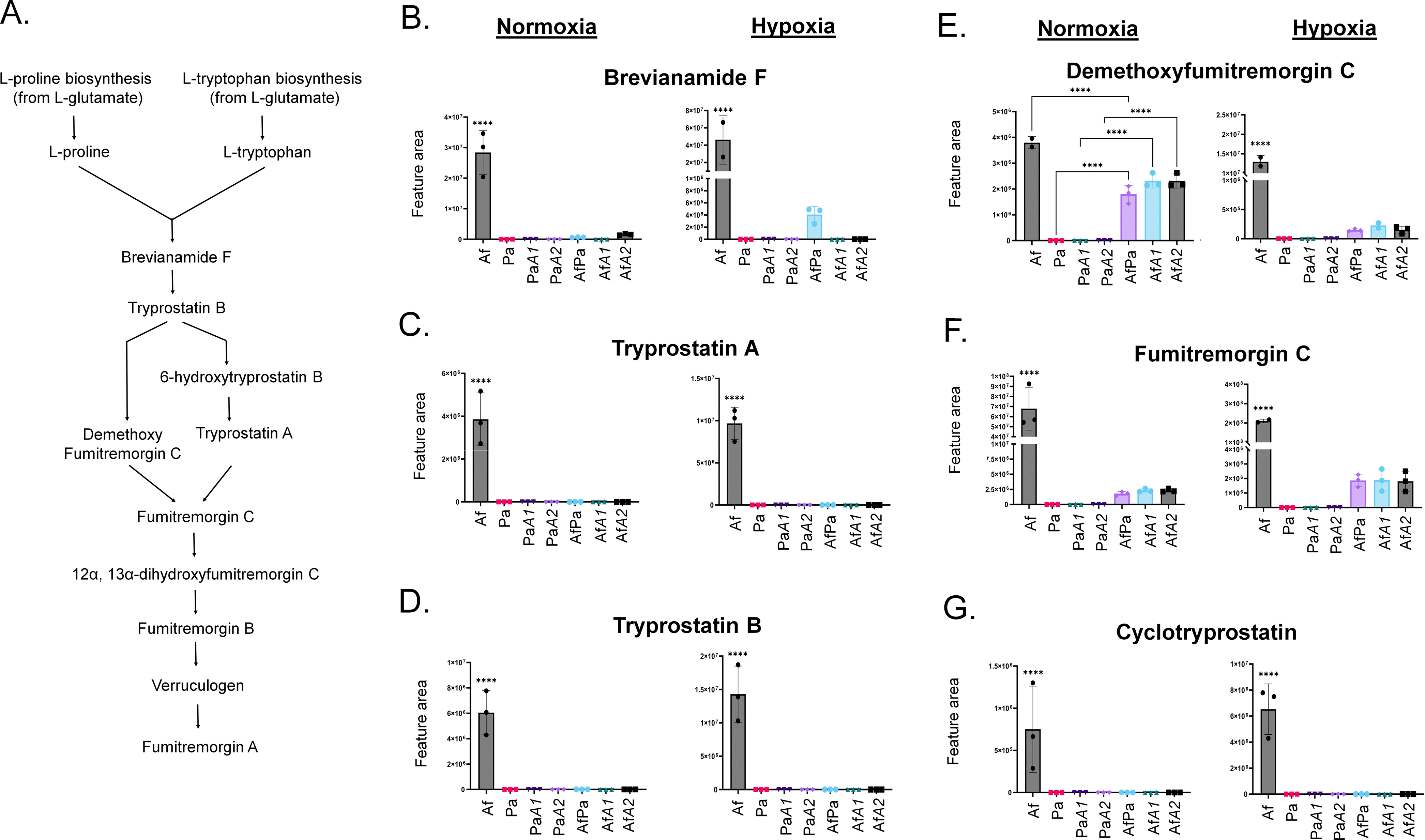
*A. fumigatus* biofilm formation induces the production of metabolites in the superpathway of fumitremorgin biosynthesis. (A) Superpathway of fumitremorgin biosynthesis. Areas of the chromatograms of (B) brevianamide F, (C) tryprostatin A, (D) tryprostatin B, (E) demethoxyfumitremorgin C, (F) fumitremorgin C, and (G) cyclotryprostatin. The results are the average of three repetitions ± standard deviation. Pa = *P. aeruginosa* wild-type; Pa*A1 = P. aeruginosa phzA1* mutant; PaA2 = *P. aeruginosa ΔphzA2* mutant; AfPa = *A. fumigatus* + *P. aeruginosa* wild-type; Af*A1 = A. fumigatus + P. aeruginosa phzA1* mutant; and AfA2 = *A. fumigatus* + *P. aeruginosa ΔphzA2* mutant.

### There is increased production of *A. fumigatus* metabolites important for iron metabolism during biofilm formation

We annotated several metabolites relevant for iron assimilation, such as ferrichrome, ferricrocin and triacetylfusigen, produced during *A. fumigatus* biofilm formation under normoxia and hypoxia conditions (Figures 2, 6A to 6C). Methyl ferrichrome is produced by Af biofilm only under normoxia (Figures 2A and 6D) while coprogen B and fusarinine B are produced only in hypoxia conditions (Figures 2B, 6E and 6F). Ferrichrome, ferricrocin, and triacetylfusigen are produced during mixed fungus-bacterial biofilm formation in both normoxia and hypoxia conditions, although in levels 10 to 1000-fold lower than in Af-only biofilms (Figures 2A, 6A, 6B, and 6C). *P. aeruginosa* wild type (Pa) and mutant strains (A1 and A2) behaved in similar patterns regarding the inhibition of production of those compounds, with a few exceptions, as AfA1 biofilms that did not produce ferrychrome in hypoxia and triacetylfusigen in normoxia.

**Figure 6.**
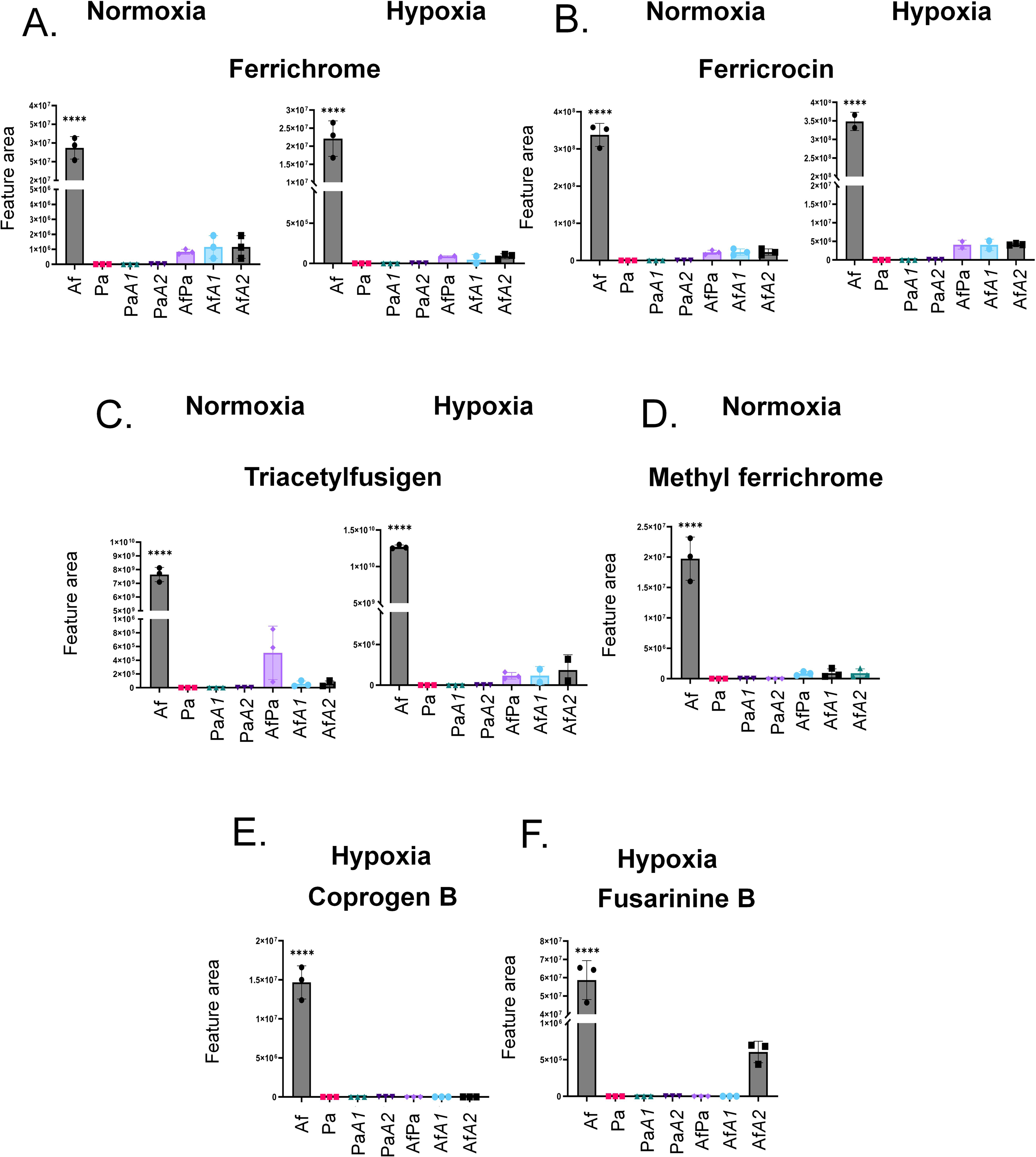
There is increased production of *A. fumigatus* metabolites important for iron metabolim during biofilm formation. Areas of the chromatograms of (A) ferrichrome, (B) ferrichrocin, (C) triacetylfusigen, (D) methylferrichrome, (E) coprogen B, and (F) fusarinine B. The results are the average of three repetitions ± standard deviation. Pa = *P. aeruginosa* wild-type; Pa*A1 = P. aeruginosa phzA1* mutant; PaA2 = *P. aeruginosa ΔphzA2* mutant; AfPa = *A. fumigatus* + *P. aeruginosa* wild-type; Af*A1 = A. fumigatus + P. aeruginosa phzA1* mutant; and AfA2 = *A. fumigatus* + *P. aeruginosa ΔphzA2* mutant.

Taken together these results strongly indicate that *A. fumigatus* can still produce several metabolites important for iron-chelation during biofilm formation in both normoxia and hypoxia conditions, notably ferrichrome, ferricrocin, and triacetylfusigen. Nevertheless, the results also show that *P. aeruginosa* strongly inhibits the overall production of all iron chelators detected in this approach, suggesting that competition for this micronutrient is a key point in *A. fumigatus*-*P. aeruginosa* interaction in biofilms.

### Gliotoxin, pyripyropene A, and fumiquinazoline F/G are produced by *A. fumigatus* during biofilm formation

Gliotoxin (GT) and GT-modified forms, such as bis(methylthio)GT, GT E, and GT G are produced during Af biofilm formation under normoxia and hypoxia conditions (Figures 2, 7A to 7D). GT production is 2-3-fold induced in AfPa, AfA1, and AfA2 during biofilm formation under normoxia conditions (Figure 2A and 7A). Curiously, under hypoxia conditions, GT levels are lower than Af in the mixed biofilms and the lack of *phzA2* function suppresses GT production completely (Figures 2B and 7B). Pyripyropene A is produced in comparable levels in all conditions, both in Af-only or mixed biofilms (Figures 2 and 7E), indicating that *P. aeruginosa* presence does not influence its production. A different pattern is seen for the production of fumiquinazoline F/G in mixed biofilms, that is completely inhibited by the bacteria in normoxia, but only partially during hypoxia (Figures 2B and 7F).

**Figure 7.**
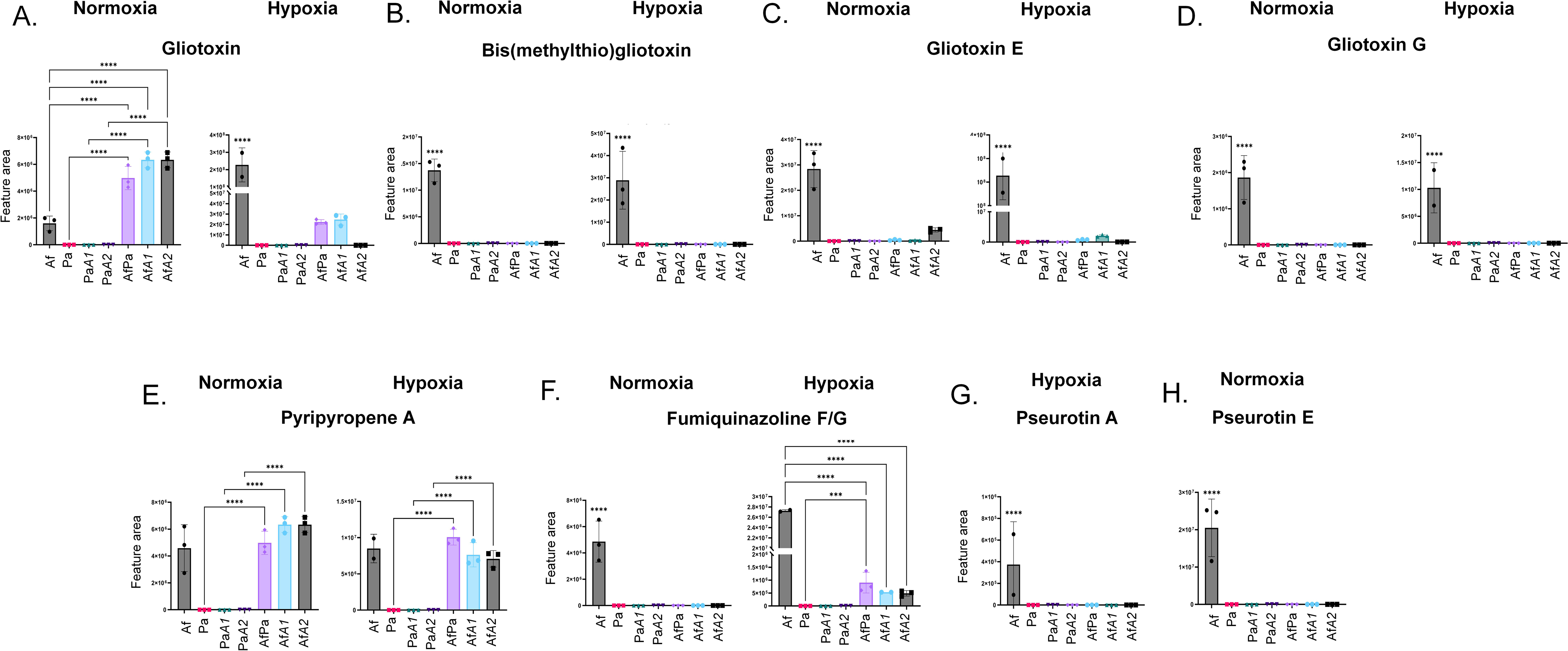
Gliotoxin, pyripyropene A, and fumiquinazoline F/G are produced during *A. fumigatus* biofilm formation. Areas of the chromatograms of (A) gliotoxin, (B) bisdethiobis(methylthio)-gliotoxin, (C) gliotoxin E, (D) gliotoxin G, (E) pyripiropene A, (F) fumiquinazoline F/G, (G) pseurotin A, and (H) pseurotin E. The results are the average of three repetitions ± standard deviation. Pa = *P. aeruginosa* wild-type; Pa*A1 = P. aeruginosa phzA1* mutant; PaA2 = *P. aeruginosa ΔphzA2* mutant; AfPa = *A. fumigatus* + *P. aeruginosa* wild-type; Af*A1 = A. fumigatus + P. aeruginosa phzA1* mutant; and AfA2 = *A. fumigatus* + *P. aeruginosa ΔphzA2* mutant.

Taken together, these results emphasize the importance of GT, pyripyropene A, and fumiquinazoline F/G during AfPa biofilm formation and suggest that *A. fumigatus* uses GT as a defense against *P. aeruginosa*.

## Discussion

The clinical and scientific interest in the co-infection with *A. fumigatus* and *P. aeruginosa* are due to its association with a decline in the lung function in cystic fibrosis patients, which has been shown in many reports (1). Inside the host, both pathogens have to face a hostile environment triggering general and specific responses and adapting to specific conditions and nutrient availability. Furthermore, they may interact with each other, which can boost their growth or lead to the production of antagonistic molecules. Among such molecules, some SM have been described affecting fungal and bacterial growth and their metabolism. However, to the best of our knowledge, an overview of the SM production by both microorganisms in co-cultivation is missing. Herein, we show that *P. aeruginosa* has antagonistic effects against *A. fumigatus* in mixed biofilms, both in normoxic and hypoxic conditions, which is in agreement with reports showing antagonistic action for bacteria isolated from clinical pulmonary samples in normal oxygen atmosphere (25, 36) and another report that showed that *P. aeruginosa* inhibitory effects are effective independently of the local oxygen pressure (37).

*P. aeruginosa* is a non-fermentative bacterium that grows anaerobically when nitrate is available, what may also be a key factor for cultivation in hypoxic conditions (37). In our work, we decided to use RPMI 1640 buffered with HEPES (pH 7.0) because it is a medium that has been used to induce biofilm formation and mimics the human plasma constitution. RPMI 1640 has an expressive amount of calcium nitrate [Ca (NO_3_)_2_ (0.42mM)], which could support *P. aeruginosa* growth in hypoxia. *P. aeruginosa* can also survive in oxygen-limited conditions using pyocyanin as an electron acceptor to regenerate NAD^+^ (38). We performed species-specific qPCR to distinguish between *A. fumigatus* and *P. aeruginosa* in co-cultured biofilms and found that *A. fumigatus* growth is inhibited by the presence of *P. aeruginosa* wild-type and *phzA1* and Δ*phzA2* mutants independently of the oxygen pressure. However, *P. aeruginosa* wild-type *and phzA1* had their growth boosted when co-cultured with *A. fumigatus* in normoxia. This result disagrees with the work performed by (39) that showed a mutually antagonistic relationship between *P. aeruginosa* and *A. fumigatus*, but confirms the data of Margalit and colleagues (40) that demonstrated that *A. fumigatus* secretome could stimulate the growth of *P. aeruginosa*. Specifically, *A. fumigatus* can produce an amino acid-rich environment in which *P. aeruginosa* can proliferate better when in co-culturing (40). The increased *P. aeruginosa* proliferation was not observed in hypoxic conditions. We hypothesize that this occurs because the fungus does not grow well in hypoxia and possibly it is not creating this amino acid-rich environment that is able to boost *P. aeruginosa* growth. Moreover, *P. aeruginosa* growth rate in hypoxia is lower than in aerobic conditions, even in the presence of nitrate and arginine, which can also be used as terminal electron acceptors (41).

The antifungal effect of *P. aeruginosa*-produced compounds on *A. fumigatus* has been extensively studied and several molecules can interfere with fungal morphology, physiology and growth. In our work, we annotated many phenazines, such as PYO, PCN, PCA, and 1-HP, that are produced by the bacterium in single and mixed cultures, both in hypoxia and normoxia. The antagonistic actions of phenazines are attributed to their redox potential since reduced phenazines are oxidized in the fungal cell by oxygen and NADPH through a NapA-dependent oxidative stress response, generating reactive oxygen species (ROS) (19; 24). All phenazines at high concentrations induce ROS and reactive nitrogen species (RNS) production by *A. fumigatus* mitochondria, which are released into the cytoplasm, leading to fungal death (42). 1-HP is the most active phenazine against *A. fumigatus* and in addition to ROS and RNS production, its high inhibitory activity is due to a specific iron chelation property (43). *P. aeruginosa* phenazine production is controlled by a complex regulatory network that involves quorum sensing and catabolite repression (44). Two redundant *phz* gene operons are responsible for phenazine-1-carboxylate production (PCA) (44, 45, 46, 47). Earlier work has shown that in *P. aeruginosa* colony biofilms, *phzA2* was expressed at high levels whereas *phzA1* is the most important operon for PCA production (45). Recent analysis suggests a dominant role of *phzA2*, resulting in a 10-fold higher expression of *phzA2* compared to *phzA1* and *phzA2* operon as the main responsible for PCA production (47), but there are marked differences in quorum-sensing regulated traits depending on the particular *P. aeruginosa* strain and specific growth conditions. Surprisingly, in contrast, our results showed that PC production is higher in Δ*phzA2* both in monoculture or *A. fumigatus* co-culture than in the *phzA1* mutant.

Another *P. aeruginosa*-produced compound that is induced upon iron starvation is pyoverdine (19), and some authors have shown that it is the main mediator of antifungal activity on *A. fumigatus* biofilms (19, 22). There are also some reports describing that pyoverdine is less produced in iron-limiting conditions (22) and in hypoxia (37). However, our results show that *P. aeruginosa* is able to produce it in RPMI medium, which is a poor-iron medium, and in low oxygen concentration atmosphere. Except for pyoverdine D, there is little influence of the *phzA1* and *phzA2* null mutations on the production of pyoverdine C, -D, -E, as expected, since the regulation of phenazines and pyoverdine are independent.

In our conditions, the rhamnolipids Rha-C10-C10, and Rha-Rha-C10-C10 were also detected in all *P. aeruginosa* strains supernatants and growth in co-culture with *A. fumigatus* increased their concentration (Figure 4 D, E). Rhamnolipids are surfactants released by *P. aeruginosa* with several roles, such as allowing swarming motility and solubilizing hydrophobic compounds that can be used as carbon and energy sources. In host-pathogens interactions, rhamnolipids are considered as virulence factors, as they may help to lyse host cells membranes, interfere with signaling pathways and solubilize the lung surfactant. They are also toxic to other bacteria, fungi and other microorganisms, conferring a competitive advantage in colonizing multiple environments (48). In mixed AfPa biofilms, the induction of rhamnolipids production might be one of the factors that interferes with *A. fumigatus* growth, but their specific role in these interactions could not be addressed in this work.

Recently, quantitative proteomic analysis showed that *A. fumigatus* exposed to *P. aeruginosa* culture filtrate had increased expression of proteins involved in SM biosynthesis, such as gliotoxin, fumagillin, and pseurotin A (49). To shed light on how *A. fumigatus* responds to all these compounds produced by *P. aeruginosa*, we also annotated and performed relative quantification of fungal SMs. We detected 20 SM secreted by *A. fumigatus* either in monoculture or in co-culture with *P. aeruginosa*. All these compounds are secreted during fungal biofilm formation either in normoxia or hypoxia. However, we were only able to detect and annotate eight compounds (demethoxyfumitremorgin C, fumitremorgin, ferrichrome, ferricrocin, traicetylfusigen, gliotoxin, gliotoxin E, and pyripyropene A) are produced during the biofilm formation by the co-culture of *A. fumigatus* and *P. aeruginosa* upon both normoxia and hypoxia conditions. Interestingly, brevianamide F and fumiquinazoline F/G are produced only upon hypoxia conditions while methyl ferrichrome only upon normoxia conditions. There results indicate that these SMs are important for the interaction between *A. fumigatus* and *P. aeruginosa*, and the production of some of them is regulated by the oxygen condition. Of note, gliotoxin was the only SM produced in higher levels in mixed biofilms as compared to Af-only biofilms, suggesting that the fungus specifically overproduces this compound in response to the bacterial antagonist. Gliotoxin (GT) has been the most well studied and characterized SM from *A. fumigatus*, and it is also important for the interaction with Pa and for biofilm formation (50, 51). Reece and colleagues (39) showed that gliotoxin has an antibacterial activity and anti-biofilm effect against several bacteria including *P. aeruginosa*. Our results emphasize the importance of gliotoxin during the interaction between *A. fumigatus* and *P. aeruginosa*, because it is the only compound produced by *A. fumigatus* in significantly higher amounts in mixed biofilms as compared to Af-only biofilms, despite the smaller fungal DNA copy number in the presence of bacteria (Figure 7A). It would be interesting to investigate the molecular mechanisms involved in such overexpression of GT induced by *P. aeruginosa.* There is little influence of *phzA1* and *phzA2* mutations on the gliotoxin production in normoxia as compared to wild type Pa. However, lack of *phzA2* function upon hypoxia decreases dramatically gliotoxin production, which might correlate with higher levels of phenazines in this condition, except for PYO (Figure 3). Further work is needed to investigate if gliotoxin has indeed a direct role in the *A. fumigatus*-*P. aeruginosa* interaction and to unravel its effects in bacterial physiology in mixed biofilms.

*P. aeruginosa* produces iron-chelators that may cause an iron-starvation environment for the fungus resulting in its anti-*Aspergillus* effect. However, the fungus counterattacks the iron-deficiency also producing siderophores (23). Siderophores are ferric-iron chelators, structurally separated in different classes named hydroxamates, catecholates, carboxylates, phenolates, and mixed-classes. Hydroxamates are sub-divided into rhodotorulic acid, ferrioxamine-, fusarinine-, coprogen-, and ferrichrome-type siderophores and are the ones that are produced by *A. fumigatus* (52). In our assay, fusarinine B, coprogen B, and ferrichrome were detected in monoculture and co-cultures. The RPMI medium is an iron-deficient medium, similar to human plasma, and that is probably the reason why *A. fumigatus* produces many siderophores in monoculture. However, only ferrichrome, ferricrocin, and triacetylfusigen are produced by *A. fumigatus* in the presence of *P. aeruginosa* during biofilm formation in both normoxia and hypoxia. Ferrichrome and ferricrocin are produced in comparable amounts in the presence of both *P. aeruginosa* wild-type and mutant strains upon normoxia or hypoxia. However, triacetylfusigen is produced in larger amounts in the presence of the *P. aeruginosa* wild type than the mutant strains. In hypoxia conditions, fusarinine B is produced only in the presence of *P. aeruginosa* Δ*phzA2* mutant, again indicating an interaction between the *phz* operons and *A. fumigatus* SM production. Previously, the importance of *A. fumigatus* siderophores for the iron competition with *P. aeruginosa* has been reported and the participation of several genetic determinants (*hapX*, *sidA*, *sidF*, *sidG* and *mirB*) involved in iron starvation adaptation in response to *P. aeruginosa* 1-HP has been demonstrated (19, 42, 43, 53). However, to the best of our knowledge that is the first time *A. fumigatus* siderophores are directly identified during the *A. fumigatus*-*P. aeruginosa* biofilm formation.

*A. fumigatus* also produced pyripyropene in mono- or dual-cultures. Pyripyropene A was originally identified as a potent inhibitor of acyl-CoA cholesterol acyltransferase (54; 55; 56), a mammalian intracellular enzyme located in the endoplasmic reticulum that forms cholesteryl esters from cholesterol (57). Pyripyropene A also shows insecticidal activity against agricultural insect pests (58; 56). It is not known if there is any correlation between these activities and aspergillosis or in the competition with bacteria, and it is a completely novel observation its identification in *A. fumigatus-P. aeruginosa* dual-cultures. In our mixed biofilm settings, the bacterial target remains to be uncovered.

Other SMs secreted by the fungus during the co-culture interaction belong to the superpathway of fumitremorgin, which are prenylated indole alkaloids compounds produced by *A. fumigatus* and *Penicillium* spp that can act as mycotoxins (59). All the compounds identified in the fumitremorgin superpathway are produced during *A. fumigatus* biofilm formation. However, only demethoxyfumitremorgin C and fumitremorgin are produced during the interaction with *P. aeruginosa* wild-type and mutant strains. Curiously, brevianamide F is produced only upon hypoxia in the presence of *P. aeruginosa* wild type but not mutant strains, what may suggest that *phzA1* and *phzA2* functions are important for brevianamide F production. Demethoxyfumitremorgin C has been shown to inhibit the cell viability and induce apoptosis of PC3 human advanced prostate cancer cells (60). Fumitremorgin C has been described as an inhibitor of a multidrug resistance protein that mediates resistance to chemotherapics in breast cancer treatment, inhibiting the growth of several phytopathogenic fungi, lethal to brine shrimp, and displaying antifeedant activity towards armyworm (61; 62; 63). We also observed fumiquinazolines, that normally accumulate in *A. fumigatus* conidia (64) secreted during both *A. fumigatus-P. aeruginosa* mono- and dual-cultures in hypoxia. It has been reported that fumiquinazoline F from *Penicillium coryphilum* has antibacterial activity against *Staphylococcus aureus* and *Micrococcus luteus* (65). It remains to be investigated if all these compounds can affect *P. aeruginosa* physiology and growth and their mechanisms of action, if any.

In conclusion, we have annotated several SMs secreted during *A. fumigatus* and *P. aeruginosa* biofilm formation, providing several opportunities to understand the interaction between these two species. Further work will concentrate on the investigation of the roles of selected compounds in both fungal and bacterial competitors, and future data may be used for the development of novel drugs for the management of chronic infections that affect cystic fibrosis patients or other immune compromised individuals.

## Methods

### *P. aeruginosa* and *A. fumigatus* strains and growth conditions

The following species and strains were used in this work: *P. aeruginosa* UCBPP-PA14 (WT, 66), *phzA1* (PA14 *phzA1::Mr7,* 67) and Δ*phzA2* (PA14 with an in-frame deletion in *phzA2,* a gift from E. Déziel), and *A. fumigatus* CEA17. *P. aeruginosa* was grown from frozen stocks (LB medium plus 20% glycerol) in solid LB for 24h at 37°C. A single colony was transferred to 30 mL of LB and cultured overnight at 37°C, 200 rpm. The culture was centrifuged at 4000 *g*, for 5 min, and the pellet was washed with 10 mL of PBS (phosphate-puffered saline). After centrifugation, the pellet was resuspended in LB and the inoculum was adjusted using a spectrophotometer to OD_600_ = 0.07-0.075. This inoculum was grown in 30 mL of LB at 37°C, 200 rpm for 5h, and the centrifugation and PBS-washing processes were repeated. The final pellet was resuspended in RPMI-Hepes and the inoculum was adjusted to OD = 0.07-0.075 (approximately 5-8 x10^8^ CFU/mL).

*A. fumigatus* strains were grown from frozen stocks and conidia suspension was obtained harvesting minimal medium plates with the grown mycelia as described by Ries and colleagues (68).

### Dual biofilm formation between *P. aeruginosa* and *A. fumigatus*

To measure the interaction between *P. aeruginosa* and *A. fumigatus*, and to determine the SM produced by them in single or co-cultures, 1 X 10^5^ CFU/mL of *P. aeruginosa* were inoculated with or without 1 X 10^6^ conidia/mL of *A. fumigatus* in 15 mL of RPMI-Hepes medium into polystyrene Petri dishes (60 x 15 mm) under hypoxic (1% O_2_, 5% CO_2_) or normoxic (approximately 20 % O_2_ and 0.04 % CO_2_) conditions, at 37°C. After 5-days, the supernatant was collected, the plate washed with 10 mL of ultrapure water to collect the cells and both were transferred to a 50 mL tube. This mixture was centrifuged at 4000 *g*, for 15 min, at 4°C to obtain the pellet, that was used for q-PCR assays, and the supernatant (20 mL), which was filtered through a 0.22 μm filter, frozen and lyophilized for SM extraction.

On the bottom of the small Petri dishes used in this experiment, biofilm production was measured by the crystal violet (CV) method. The biofilm was dried at 37°C for 30 min and then stained with 5 mL of 0.05% (w/v) CV for 10 min. The plates were washed with 50 mL of PBS and the CV was solubilized with 3 mL of 95% ethanol. 100 uL samples were transferred to 96-well plates and the absorbance at 595 nm was determined, as a measure for biofilm formation.

### Dual Quantification of Species-Specific Biofilm Growth by qPCR

For DNA extraction, the pellet obtained from *P. aeruginosa* - *A. fumigatus* interaction was frozen and lyophilized, before being triturating by adding 2 mm glass beads and 0.1 mm zirconia/silica beads and vortexed for 5 minutes. To the resulting powder, 1 mL of extraction buffer (EB) was added and the tubes vortexed for 5 min. Tubes were incubated in a water bath at 70°C for 45 min, (every 10-15 min, the tubes were removed from the water bath and 5 min-vortexed). One milliliter of phenol:chloroform (1:1) was added to the mixture and vortexed for more 5 min. The content was transferred to 2 mL tubes and centrifuged at 14,000g for 15 min at room temperature. The supernatant was collected and transferred to 1.5 mL tubes, and 600 μL isopropanol (MERCK S. A) were added. Samples were incubated at 4°C for 1h before being centrifuged at 14,000 *g* for 15 min at 4°C. The supernatant was discarded, the pellet washed with 200 μL 70% ethanol, air dried for 15 min at room temperature and the pellet was resuspended in deionized water and treated with 50 RNAse (Promega).

*P. aeruginosa* DNA was specifically quantified by qPCR with primers for the *ecfX* (ecfX-F, 5’-CGCATGCCTATCAGGCGTT-3’ and ecfX-R, 5’-GAACTGCCCAGGTGCTTGC-3’) and *gyrB* (gyrB-F, 5’-CCTGACCATCCGTCGCCACAAC-3’ and gyrB-R, 5’-CGCAGCAGGATGCCGACGCC-3’) genes (31, 33). *A. fumigatus* was quantified by amplification of 18S rDNA (18S fw, 5’-GACCTCGGCCCTTAAATAGC-3’ and 18S rv, 5’-CTCGGCCAAGGTGATGTACT-3’). The 10 μL qPCR reaction was composed by 5 μL SYBR green PCR master mix kit (Applied Biosystems, Foster City, CA, USA), 2,5 pmol of each primer and 100 ng DNA. Cycling was performed on the ABI 7500 Fast real-time PCR system with an initial hold at 95°C for 15 min, followed by 45 cycles at 95°C for 15 s, and 60°C for 1 min, with a C_T_ of 35 being the threshold. Negative controls without DNA were included in each qPCR run.

### SM extraction and UHPLC-HRMS^2^ analysis

SMs were extracted from 50 mg freeze-dried samples by resuspension in 1 mL of HPLC-grade MeOH followed by 1 h of sonication in an ultrasonic bath. For sample preparation, 500 µL of the obtained extracts were 0.22 μm filtered, transferred to vials and diluted with HPLC -grade MeOH to a total volume of 1 mL.

UHPLC-HRMS^2^ positive mode analysis was performed in a Thermo Scientific QExactive Hybrid Quadrupole-Orbitrap Mass Spectrometer coupled to a Dionex UltiMate 3000 RSLCnano UHPLC System. As stationary phase, a Thermo Scientific column Accucore C18 2.6 µm (2.1 mm × 100 mm) was used. Mobile phase was 0.1% formic acid (A) and acetonitrile + 0.1% formic acid (B). Eluent profile (A/B %): 95/5 up to 2/98 within 10 min, maintaining 2/98 for 5 min, down to 95/5 within 1.2 min and maintaining for 8.8 min. Total run time was 20 min for each run and flow rate of 0.3 mL.min^-1^. Injection volume was 5 µL. MS spectra were acquired with *m/*z ranges from 100 to 1500, with 70000 mass resolution. Ionization parameters: sheath gas flow rate (45), aux gas flow rate (10), sweep gas flow rate (2), spray voltage (3.5 kV), capillary temperature (250° C), S-lens RF level (50) e auxiliary gas heater temperature (400° C). MS^2^ spectra were acquired in data-dependent acquisition (DDA) mode. Normalized collision energy was applied stepwise (20, 30 and 40), and 5 most intense precursors per cycle were measured with 17500 resolution.

### UHPLC-HRMS^2^ data processing and Feature-Based Molecular Networking (FBMN)

Raw UHPLC-HRMS^2^ data were converted into mzXML format files in MSConvert (69), with 32-bit binary encoding precision, zlib compression, and peak peaking. Feature detection was performed in MZmine2 (v.2.53) (70). For MS1 spectra mass detection, an intensity threshold of 1E5 was used, and for MS2 an intensity threshold of 1E3 was used. For MS1 chromatogram building (71), a 5-ppm mass accuracy and a minimum peak intensity of 5E5 was set. Extracted ion chromatograms (XICs) were deconvolved using the baseline cut-off algorithm at an intensity of 1E5, minimum peak height of 3E5 and a peak duration range from 0.05 to 2 minutes. After chromatographic deconvolution, XICs were matched to MS2 spectra within 0.02 m/z and 0.2-minute retention time windows. Isotope peaks were grouped with 5 ppm mass tolerance, 0.1- minute retention time tolerance and a maximum charge of 2. Detected peaks in different samples were aligned with a 5-ppm tolerance, 75% weight for *m/z* and 25% for retention time. MS1 features without MS2 features assigned were filtered out the resulting matrix as well as features that did not contain isotope peaks and that did not occur in at least three samples. Finally, the feature table was exported as a .csv file, and corresponding MS2 spectra exported as .mgf files. Features observed in blank samples were filtered.

A molecular network was created with the Feature-Based Molecular Networking (FBMN) workflow (72) on GNPS (https://gnps.ucsd.edu) (73). The data was filtered by removing all MS2 fragment ions within +/- 17 Da of the precursor m/z. MS2 spectra were window filtered by choosing only the top 6 fragment ions in the +/- 50 Da window throughout the spectrum. The precursor ion mass tolerance was set to 0.02 Da and the MS2fragment ion tolerance to

1.2 Da. A molecular network was then created where edges were filtered to have a cosine score above 0.65 and more than 4 matched peaks. Further, edges between two nodes were kept in the network if and only if each of the nodes appeared in each other’s respective top 10 most similar nodes. Finally, the maximum size of a molecular family was set to 100, and the lowest scoring edges were removed from molecular families until the molecular family size was below this threshold. The spectra in the network were then searched against GNPS spectral libraries (73; 74). The library spectra were filtered in the same manner as the input data. All matches kept between network spectra and library spectra were required to have a score above 0.65 and at least 4 matched peaks. DEREPLICATOR PLUS was used to annotate MS/MS spectra (75). The molecular networks were visualized using Cytoscape software (76). Resulting networks were displayed and analyzed with Cytoscape (v.3.8.2).

### Metabolite annotation

For SM dereplication, metabolites were annotated based on the GNPS MS^2^ database via the Feature-Based Molecular Networking (FBMN) and DEREPLICATOR PLUS workflows available on the GNPS platform. Other metabolites were manually searched against natural products databases such as the Dictionary of Natural Products and the acquired MS^2^ spectra was compared to spectra deposited either on the GNPS database or previously published in the literature. Compounds Gliotoxin, Gliotoxin E, Pseurotin E and Spirotryprostatin A were annotated based on their exact masses.

### Data Availability

The molecular networking job can be publicly accessed at https://gnps.ucsd.edu/ProteoSAFe/status.jsp?task=0b027bfab9064518945341d 3c24ba2a9 and the DEREPLICATOR PLUS job at https://gnps.ucsd.edu/ProteoSAFe/status.jsp?task=415264520a41492a9b43b4 7e623028ed for hypoxia conditions and https://gnps.ucsd.edu/ProteoSAFe/status.jsp?task=08a9ebb5609745d5a6f8058 96e527b67and https://gnps.ucsd.edu/ProteoSAFe/status.jsp?task=e59e11f719cf4ae5beb4060 90a987f01for normoxia, respectively.

## Acknowledgements

We thank Fundação de Amparo à Pesquisa do Estado de São Paulo (FAPESP) 2017/19821-5 (RWB), 2017/07536-4 (ACC), 2016/12948-7 (PAC) 2016/07870-9 (GHG), 2021/07038-0 (DA), 2021/00728-0 (TF) and 2021/11062-3 (RLB), and Conselho Nacional de Desenvolvimento Científico e Tecnológico (CNPq) 301058/2019-9 and 404735/2018-5 (GHG), both from Brazil, and National Institutes of Health/National Institute of Allergy and Infectious Diseases (R01AI153356), from the USA. This study was financed in part by the Coordenação de Aperfeiçoamento de Pessoal de Nível Superior – Brasil (CAPES) – Finance Code 001 (RSL)

## References

1. Beswick E, Amich J, Gago S. 2020. Factoring in the Complexity of the Cystic Fibrosis Lung to Understand *Aspergillus fumigatus* and *Pseudomonas aeruginosa* Interactions. Pathogens. 6;9(8):639. doi: 10.3390/pathogens9080639. PMID: 32781694; PMCID: PMC7460534.

2. Vilaplana L, Marco MP. 2020. Phenazines as potential biomarkers of *Pseudomonas aeruginosa* infections: synthesis regulation, pathogenesis and analytical methods for their detection. Anal Bioanal Chem. 412(24):5897–5912. doi: 10.1007/s00216-020-02696-4. Epub 2020 May 27. PMID: 32462363.

3. Haq IJ, Gardner A, Brodlie M. 2016. A multifunctional bispecific antibody against *Pseudomonas aeruginosa* as a potential therapeutic strategy. Ann Transl Med. 4(1):12. doi: 10.3978/j.issn.2305-5839.2015.10.10. PMID: 26855948; PMCID: PMC4716945.

4. Worlitzsch D, Tarran R, Ulrich M, Schwab U, Cekici A, Meyer KC, Birrer P, Bellon G, Berger J, Weiss T, Botzenhart K, Yankaskas JR, Randell S, Boucher RC, Döring G. 2002. Effects of reduced mucus oxygen concentration in airway *Pseudomonas* infections of cystic fibrosis patients. J Clin Invest 109(3):317–25. doi: 10.1172/JCI13870. PMID: 11827991; PMCID: PMC150856.

5. Warrier A, Satyamoorthy K, Murali TS. Quorum-sensing regulation of virulence factors in bacterial biofilm. Future Microbiol. 2021 Sep;16:1003–1021. doi: 10.2217/fmb-2020-0301. Epub 2021 Aug 20. PMID: 34414776.

6. Olivares E, Badel-Berchoux S, Provot C, Prévost G, Bernardi T, Jehl F. Clinical Impact of Antibiotics for the Treatment of *Pseudomonas aeruginosa* Biofilm Infections. Front Microbiol. 2020 Jan 9;10:2894. doi: 10.3389/fmicb.2019.02894. PMID: 31998248; PMCID: PMC6962142.

7. King J, Brunel SF, Warris A. 2016. *Aspergillus* infections in cystic fibrosis. J Infect. 5;72 Suppl:S50-5. doi: 10.1016/j.jinf.2016.04.022. Epub 2016 May 11. PMID: 27177733.

8. Bastos RW, Rossato L, Goldman GH, Santos DA. 2021. Fungicide effects on human fungal pathogens: Cross-resistance to medical drugs and beyond. PLoS Pathog. 9;17(12):e1010073. doi: 10.1371/journal.ppat.1010073. PMID: 34882756; PMCID: PMC8659312.

9. Burks C, Darby A, Gómez Londoño L, Momany M, Brewer MT. 2021. Azole-resistant *Aspergillus fumigatus* in the environment: Identifying key reservoirs and hotspots of antifungal resistance. PLoS Pathog. 29;17(7):e1009711. doi: 10.1371/journal.ppat.1009711. PMID: 34324607; PMCID: PMC8321103.

10. Latgé JP, Chamilos G. *Aspergillus fumigatus* and Aspergillosis in 2019. 2019. Clin Microbiol Rev. 13;33(1):e00140-18. doi: 10.1128/CMR.00140-18. PMID: 31722890; PMCID: PMC6860006.

11. Baxter CG, Dunn G, Jones AM, Webb K, Gore R, Richardson MD, Denning DW. 2013. Novel immunologic classification of aspergillosis in adult cystic fibrosis. J Allergy Clin Immunol. 132(3):560–566.e10. doi: 10.1016/j.jaci.2013.04.007. Epub 2013 May 29. PMID: 23726262.

12. Moss A, Juarez-Colunga E, Nathoo F, Wagner B, Sagel S. 2016. A comparison of change point models with application to longitudinal lung function measurements in children with cystic fibrosis. Stat Med. 30;35(12):2058-73. doi: 10.1002/sim.6845. Epub 2016 Jan 5. PMID: 27118629.

13. Agarwal R, Sehgal IS, Dhooria S, Aggarwal AN. 2016. Developments in the diagnosis and treatment of allergic bronchopulmonary aspergillosis. Expert Rev Respir Med. 10(12):1317–1334. doi: 10.1080/17476348.2016.1249853. Epub 2016 Nov 7. PMID: 27744712.

14. Brandt C, Roehmel J, Rickerts V, Melichar V, Niemann N, Schwarz C. 2018. *Aspergillus* Bronchitis in Patients with Cystic Fibrosis. Mycopathologia. 183(1):61–69. doi: 10.1007/s11046-017-0190-0. Epub 2017 Aug 17. PMID: 28819878.

15. Bakare N, Rickerts V, Bargon J, Just-Nübling G. 2003. Prevalence of *Aspergillus fumigatus* and other fungal species in the sputum of adult patients with cystic fibrosis. Mycoses. 46(1-2):19–23. doi: 10.1046/j.1439-0507.2003.00830.x. PMID: 12588478.

16. Amin R, Dupuis A, Aaron SD, Ratjen F. 2010. The effect of chronic infection with *Aspergillus fumigatus* on lung function and hospitalization in patients with cystic fibrosis. Chest 137 171–176. 10.1378/chest.09-1103

17. Paugam A, Baixench MT, Demazes-Dufeu N, Burgel PR, Sauter E, Kanaan R, et al. 2010. Characteristics and consequences of airway colonisation by filamentous fungi in 201 adult patients with cystic fibrosis in France. Med. Mycol. 48(Suppl. 1)S32–S36. 10.3109/13693786.2010.503665

18. Hector A., Kirn T., Ralhan A., Graepler-Mainka U., Berenbrinker S., Riethmueller J., et al. (2016). Microbial colonization and lung function in adolescents with cystic fibrosis. J. Cyst. Fibros. 15 340–349. 10.1016/j.jcf.2016.01.004

19. Briard B, Mislin GLA, Latgé JP, Beauvais A (2019) Interactions between *Aspergillus fumigatus* and pulmonary bacteria: current state of the ’ield, new data, and future perspective. J Fungi 5:48. doi: 10.3390/jof5020048

20. Briard B., Rasoldier V., Bomme P., ElAouad N., Guerreiro C., Chassagne P., et al. (2017). Dirhamnolipids secreted from *Pseudomonas aeruginosa* modify anjpegungal susceptibility of *Aspergillus fumigatus* by inhibiting beta1,3 glucan synthase activity. ISME J. 11 1578–1591. 10.1038/ismej.2017.32

21. Mowat E., Rajendran R., Williams C., McCulloch E., Jones B., Lang S., et al. (2010). *Pseudomonas aeruginosa* and their small diffusible extracellular molecules inhibit *Aspergillus fumigatus* biofilm formation. FEMS Microbiol. Lett. 313 96–102. 10.1111/j.1574-6968.2010.02130.x

22. Sass G., Nazik H., Penner J., Shah H., Ansari S. R., Clemons K. V., et al. (2018). Studies of *Pseudomonas aeruginosa* mutants indicate pyoverdine as the central factor in inhibition of *Aspergillus fumigatus* biofilm. J. Bacteriol. 200:e00345–17. 10.1128/JB.00345-17

23. Sass G, Ansari SR, Dietl AM, et al (2019) Intermicrobial interaction: *Aspergillus fumigatus* siderophores protect against competition by Pseudomonas aeruginosa. PLoS One 14:e0216085. doi: 10.1371/journal.pone.0216085

24. Zheng H, Kim J, Liew M, et al (2015) Redox metabolites signal polymicrobial biofilm development via the napa oxidative stress cascade in *Aspergillus*. Curr Biol 25:29–37. doi: 10.1016/j.cub.2014.11.018.

25. Kerr JR, Taylor GW, Rutman A, Hoiby N, Cole PJ, Wilson R 1999. *Pseudomonas aeruginosa* pyocyanin and 1-hydroxyphenazine inhibit fungal growth. J. Clin. Pathol. 52 385–387. 10.1136/jcp.52.5.385.

26. Kumar SN, Nisha G, Sudaresan A, Venugopal V, Kumar MS, Lankalapalli R, et al. 2014. Synergistic Activity of Phenazines Isolated From *Pseudomonas aeruginosa* in Combination With Azoles Against Candida Species. Med. Mycol. 52, 482–490. doi: 10.1093/MMY/MYU012

27. Sass G, Nazik H, Chatterjee P, Stevens D. A. 2021. Under Nonlimiting Iron Conditions Pyocyanin Is a Major Antifungal Molecule, and Differences Between Prototypic *Pseudomonas aeruginosa* Strains. Med. Mycol. 59, 453– 464. doi: 10.1093/mmy/myaa066

28. Ramos AN, Peral MC, Valdez JC. 2010. Differences between *Pseudomonas aeruginosa* in a clinical sample and in a colony isolated from it: comparison of virulence capacity and susceptibility of biofilm to inhibitors. Comp Immunol Microbiol Infect Dis 33(3):267–75. doi: 10.1016/j.cimid.2008.10.004. Epub 2008 Nov 22. PMID: 19027954.

29. Du X, Li Y, Zhou Q, Xu Y. Regulation of gene expression in *Pseudomonas aeruginosa* M18 by phenazine-1-carboxylic acid. Appl Microbiol Biotechnol. 2015 Jan;99(2):813–25. doi: 10.1007/s00253-014-6101-0. Epub 2014 Oct 11. PMID: 25304879.

30. Lau GW, Hassett DJ, Ran H, Kong F. 2004. The role of pyocyanin in *Pseudomonas aeruginosa* infection. Trends Mol Med 10(12):599–606. doi: 10.1016/j.molmed.2004.10.002. PMID: 15567330.

31. Qin X, Emerson J, Stapp J, Stapp L, Abe P, Burns JL. 2003. Use of realtime PCR with multiple targets to identify Pseudomonas aeruginosa and other nonfermenting gram-negative bacilli from patients with cystic fibrosis. J. Clin. Microbiol 41, 4312–4317. doi: 10.1128/JCM.41.9.4312.

32. Lavenir R, Jocktane D, Laurent F, Nazaret S, Cournoyer B. 2007. Improved reliability of *Pseudomonas aeruginosa* PCR detection by the use of the species-specific *ecfX* gene target. J Microbiol Methods 70(1):20–9. doi: 10.1016/j.mimet.2007.03.008. Epub 2007 Mar 30. PMID: 17490767.

33. Anuj SN, Whiley DM, Kidd TJ, Bell SC, Wainwright CE, Nissen MD, et al. 2009. Identification of *Pseudomonas aeruginosa* by a duplex real-time polymerase chain reaction assay targeting the *ecfX* and the *gyrB* genes. Diagn. Microbiol. Infect. Dis. 63, 127–131. doi: 10.1016/j.diagmicrobio.2008.09.018.

34. Barakat R, Goubet I, Manon S, Berges T, Rosenfeld E. Unsuspected pyocyanin effect in yeast under anaerobiosis. Microbiologyopen. 2014 Feb;3(1):1–14. doi: 10.1002/mbo3.142. Epub 2013 Dec 5. PMID: 24307284; PMCID: PMC3937724.

35. Steffan N, Grundmann A, Yin WB, Kremer A, Li SM. 2009. Indole prenyltransferases from fungi: a new enzyme group with high potential for the production of prenylated indole derivatives. Curr Med Chem. 16(2):218–31. doi: 10.2174/092986709787002772. PMID: 19149573.

36. Yadav V Gupta J Mandhan R Chhillar AK Dabur R Singh DD Sharma GL 2005. Investigations on anti-Aspergillus properties of bacterial products. Lett Appl Microbiol 41: 309–314.

37. Anand R, Clemons KV, Stevens DA. Effect of Anaerobiasis or Hypoxia on *Pseudomonas aeruginosa* Inhibition of *Aspergillus fumigatus* Biofilm. Arch Microbiol. 2017 Aug;199(6):881–890. doi: 10.1007/s00203-017-1362-5. Epub 2017 Mar 29. PMID: 28357473.

38. Price-Whelan A, Dietrich LE, Newman DK. 2007. Pyocyanin alters redox homeostasis and carbon flux through central metabolic pathways in *Pseudomonas aeruginosa* PA14. J Bacteriol. 189(17):6372–81. doi: 10.1128/JB.00505-07. Epub 2007 May 25. PMID: 17526704; PMCID: PMC1951912.

39. Reece E, Doyle S, Greally P, Renwick J, McClean S. 2018. *Aspergillus fumigatus* Inhibits *Pseudomonas aeruginosa* in co-culture: implications of a mutually antagonistic relationship on virulence and inflammation in the CF airway. Front. Microbiol. 9:1205. 10.3389/fmicb.2018.01205

40. Margalit A, Carolan JC, Sheehan D, Kavanagh K. 2020. The *Aspergillus fumigatus* Secretome Alters the Proteome of *Pseudomonas aeruginosa* to Stimulate Bacterial Growth: Implications for Co-infection. Mol Cell Proteomics. 19(8):1346–1359. doi: 10.1074/mcp.RA120.002059. Epub 2020 May 23. PMID: 32447284; PMCID: PMC8015003.

41. Filiatrault MJ, Picardo KF, Ngai H, Passador L, Iglewski BH. 2006. Identification of *Pseudomonas aeruginosa* genes involved in virulence and anaerobic growth. Infect Immun. 74(7):4237–45. doi: 10.1128/IAI.02014-05. PMID: 16790798; PMCID: PMC1489737.

42. Moree WJ, Phelan VV, Wu CH, Bandeira N, Cornett DS, Duggan BM, Dorrestein PC. Interkingdom metabolic transformations captured by microbial imaging mass spectrometry. Proc Natl Acad Sci U S A. 2012 Aug 21;109(34):13811–6. doi: 10.1073/pnas.1206855109. Epub 2012 Aug 6. PMID: 22869730; PMCID: PMC3427086.

43. Briard B, Bomme P, Lechner BE, Mislin GLA, Lair V, Prévost M-C et al. 2015. *Pseudomonas aeruginosa* manipulates redox and iron homeostasis of its microbiota partner *Aspergillus fumigatus* via phenazines. Sci Rep 5: 8220.

44. Sultan M, Arya R, Kim KK. 2021. Roles of Two-Component Systems in *Pseudomonas aeruginosa* Virulence. Int J Mol Sci. 10;22(22):12152. doi: 10.3390/ijms222212152. PMID: 34830033; PMCID: PMC8623646.

45. Recinos DA, Sekedat MD, Hernandez A, Cohen TS, Sakhtah H, Prince AS, Price-Whelan A, Dietrich LE. 2021. Redundant phenazine operons in *Pseudomonas aeruginosa* exhibit environment-dependent expression and differential roles in pathogenicity. Proc Natl Acad Sci U S A. 20;109(47):19420-5. doi: 10.1073/pnas.1213901109. Epub 2012 Nov 5. PMID: 23129634; PMCID: PMC3511076.

46. Cui Q, Lv H, Qi Z, Jiang B, Xiao B, Liu L, Ge Y, Hu X. 2016. Cross-Regulation between the phz1 and phz2 Operons Maintain a Balanced Level of Phenazine Biosynthesis in *Pseudomonas aeruginosa* PAO1. PLoS One. 6;11(1):e0144447. doi: 10.1371/journal.pone.0144447. PMID: 26735915; PMCID: PMC4703396.

47. Schmitz S, Rosenbaum MA. 2020. Controlling the Production of *Pseudomonas* Phenazines by Modulating the Genetic Repertoire. ACS Chem Biol. 18;15(12):3244-3252. doi: 10.1021/acschembio.0c00805. Epub 2020 Dec 1. PMID: 33258592.

48. Soberón-Chávez G, Lépine F, Déziel E. 2005. Production of rhamnolipids by *Pseudomonas aeruginosa*. Appl Microbiol Biotechnol. 68(6):718–25. doi: 10.1007/s00253-005-0150-3. Epub 2005 Oct 13. PMID: 16160828.

49. Margalit A, Sheehan D, Carolan JC, Kavanagh K. 2022. Exposure to the *Pseudomonas aeruginosa* secretome alters the proteome and secondary metabolite production of *Aspergillus fumigatus*. Microbiology (Reading). 168(3). doi: 10.1099/mic.0.001164. PMID: 35333152.

50. Bruns S, Seidler M, Albrecht D, Salvenmoser S, Remme N, Hertweck C, Brakhage AA, Kniemeyer O, Müller FM. Functional genomic profiling of *Aspergillus fumigatus* biofilm reveals enhanced production of the mycotoxin gliotoxin. Proteomics. 2010 Sep;10(17):3097–107. doi: 10.1002/pmic.201000129. PMID: 20645385.

51. Dolan SK, O’Keeffe G, Jones GW, Doyle S. 2015. Resistance is not futile: gliotoxin biosynthesis, functionality and utility. Trends Microbiol. 23(7):419–28. doi: 10.1016/j.tim.2015.02.005. Epub 2015 Mar 10. PMID: 25766143.

52. Aguiar M, Orasch T, Misslinger M, Dietl AM, Gsaller F, Haas H. The Siderophore Transporters Sit1 and Sit2 Are Essential for Utilization of Ferrichrome-, Ferrioxamine- and Coprogen-Type Siderophores in *Aspergillus fumigatus*. J Fungi (Basel). 2021 Sep 16;7(9):768. doi: 10.3390/jof7090768. PMID: 34575806; PMCID: PMC8470733.

53. Nazik H, Sass G, Ansari SR, Ertekin R, Haas H, Déziel E, Stevens DA. 2020. Novel intermicrobial molecular interaction: *Pseudomonas aeruginosa* Quinolone Signal (PQS) modulates *Aspergillus fumigatus* response to iron. Microbiology (Reading). 166(1):44–55. doi: 10.1099/mic.0.000858. Epub 2019 Nov 15. PMID: 31778108.

54. Omura S, Tomoda H, Kim YK, Nishida H. 1993. Pyripyropenes, highly potent inhibitors of acyl-CoA:cholesterol acyltransferase produced by *Aspergillus fumigatus*. J Antibiot (Tokyo) 46(7):1168–9. doi: 10.7164/antibiotics.46.1168. PMID: 8360113.

55. Tomoda H, Kim YK, Nishida H, Masuma R, Omura S. 1994. Pyripyropenes, novel inhibitors of acyl-CoA:cholesterol acyltransferase produced by *Aspergillus fumigatus*. I. Production, isolation, and biological properties. J Antibiot (Tokyo) 47(2):148–53. doi: 10.7164/antibiotics.47.148. PMID: 8150709.

56. Raffa N, Keller NP. 2019. A call to arms: Mustering secondary metabolites for success and survival of an opportunistic pathogen. PLoS Pathog. 4;15(4):e1007606. doi: 10.1371/journal.ppat.1007606. PMID: 30947302; PMCID: PMC6448812.

57. Spector AA, Mathur SN, Kaduce TL. 1979. Role of acylcoenzyme A: cholesterol o-acyltransferase in cholesterol metabolism. Prog. Lipid Res. 18 (1): 31–53. doi:10.1016/0163-7827(79)90003-1.

58. Goto K, Horikoshi R, Nakamura S, Mitomi M, Oyama K, Hirose T, Sunazuka T, Ōmura S. 2019. Synthesis of pyripyropene derivatives and their pest-control efficacy. J Pestic Sci. 25;44(4):255-263. doi: 10.1584/jpestics.D19-032. PMID: 31777444; PMCID: PMC6861429.

59. Maiya S, Grundmann A, Li SM, Turner G. 2006. The fumitremorgin gene cluster of *Aspergillus fumigatus*: identification of a gene encoding brevianamide F synthetase. Chembiochem. 7(7):1062–9. doi: 10.1002/cbic.200600003. PMID: 16755625.

60. Kim YS, Kim SK, Park SJ. 2017. Apoptotic effect of demethoxyfumitremorgin C from marine fungus *Aspergillus fumigatus* on PC3 human prostate cancer cells. Chem Biol Interact. 1;269:18–24. doi: 10.1016/j.cbi.2017.03.015. Epub 2017 Mar 28. PMID: 28359723.

61. Rabindran SK, Ross DD, Doyle LA, Yang W, Greenberger LM. 2000. Fumitremorgin C reverses multidrug resistance in cells transfected with the breast cancer resistance protein. Cancer Res. 1;60(1):47-50. PMID: 10646850.

62. González-Lobato L, Real R, Prieto JG, Alvarez AI, Merino G. 2010. Differential inhibition of murine Bcrp1/Abcg2 and human BCRP/ABCG2 by the mycotoxin fumitremorgin C. Eur J Pharmacol. 10;644(1-3):41-8. doi: 10.1016/j.ejphar.2010.07.016. Epub 2010 Jul 22. PMID: 20655304.

63. Li XJ, Zhang Q, Zhang AL, Gao JM. 2012. Metabolites from *Aspergillus fumigatus*, an endophytic fungus associated with *Melia azedarach*, and their antifungal, antifeedant, and toxic activities. J Agric Food Chem. 4;60(13):3424-31. doi: 10.1021/jf300146n. Epub 2012 Mar 23. PMID: 22409377.

64. Lim FY, Ames B, Walsh CT, Keller NP. 2014. Co-ordination between BrlA regulation and secretion of the oxidoreductase FmqD directs selective accumulation of fumiquinazoline C to conidial tissues in *Aspergillus fumigatus*. Cell Microbiol. 16(8):1267–83. doi: 10.1111/cmi.12284. Epub 2014 Apr 15. PMID: 24612080; PMCID: PMC4114987.

65. Silva MG, Furtado NA, Pupo MT, Fonseca MJ, Said S, da Silva Filho AA, Bastos JK. 2004. Antibacterial activity from Penicillium corylophilum Dierckx. Microbiol Res. 159(4):317–22. doi: 10.1016/j.micres.2004.06.003. PMID: 15646377.

66. Rahme LG, Stevens EJ, Wolfort SF, Shao J, Tompkins RG, Ausubel FM. 1995. Common virulence factors for bacterial pathogenicity in plants and animals. Science. 30;268(5219):1899-902. doi: 10.1126/science.7604262. PMID: 7604262.

67. Liberati NT, Urbach JM, Miyata S, Lee DG, Drenkard E, Wu G, Villanueva J, Wei T, Ausubel FM. 2006. An ordered, nonredundant library of *Pseudomonas aeruginosa* strain PA14 transposon insertion mutants. Proc Natl Acad Sci U S A. 21;103(8):2833-8. doi: 10.1073/pnas.0511100103. Epub 2006 Feb 13. Erratum in: Proc Natl Acad Sci U S A. 2006 Dec 26;103(52):19931. PMID: 16477005; PMCID: PMC1413827.

68. Ries LNA, Beattie SR, Espeso EA, Cramer RA, Goldman GH. 2016. Diverse regulation of the CreA carbon catabolite repressor in *Aspergillus nidulans*. Genetics 203 335–352. 10.1534/genetics.116.187872

69. Chambers MC, Maclean B, Burke R, Amodei D, Ruderman DL, Neumann S, et al. 2012. A cross-platform toolkit for mass spectrometry and proteomics. Nature Biotechnology. 30(10):918–20.

70. Pluskal T, Castillo S, Villar-Briones A, Orešič M. 2010. MZmine 2: Modular framework for processing, visualizing, and analyzing mass spectrometry-based molecular profile data. BMC Bioinformatics.11(1):395.

71. Myers OD, Sumner SJ, Li S, Barnes S, Du X. 2017. One Step Forward for Reducing False Positive and False Negative Compound Identifications from Mass Spectrometry Metabolomics Data: New Algorithms for Constructing Extracted Ion Chromatograms and Detecting Chromatographic Peaks. Analytical Chemistry. 89(17):8696–703.

72. Nothias L-F, Petras D, Schmid R, Dührkop K, Rainer J, Sarvepalli A, et al. 2020. Feature-based molecular networking in the GNPS analysis environment. Nature Methods. 17(9):905–8.

73. Wang M, Carver JJ, Phelan VV, Sanchez LM, Garg N, Peng Y, et al. 2016. Sharing and community curation of mass spectrometry data with Global Natural Products Social Molecular Networking. Nature biotechnology. 34(8):828–37.

74. Horai H, Arita M, Kanaya S, Nihei Y, Ikeda T, Suwa K, et al. 2010. MassBank: a public repository for sharing mass spectral data for life sciences. Journal of mass spectrometry. 45(7):703–14.

75. Mohimani H, Gurevich A, Shlemov A, Mikheenko A, Korobeynikov A, Cao L, et al. 2018. Dereplication of microbial metabolites through database search of mass spectra. Nature Communications. 9(1):4035.

76. Shannon P, Markiel A, Ozier O, Baliga NS, Wang JT, Ramage D, et al. 2003. Cytoscape: a software environment for integrated models of biomolecular interaction networks. Genome Res. 13(11):2498–504.

